# Higher Lipid Saturation in Well-Irrigated Georgia Cotton Plants: A Field-Based NMR Metabolomics Study

**DOI:** 10.64898/2026.06.08.730374

**Authors:** Krishna Patel, Christopher Esselman, Jordan Croy, Matthew Gillis, Pedro Augusto Da Pos Rodrigues, Alvin Simmons, Ricardo M Borges, Arthur S. Edison, William E Snyder

## Abstract

Cotton (*Gossypium hirsutum*) is an economically important crop, but it faces increasing pest outbreaks, especially in non-irrigated areas. In this study, 20 cotton farms using center-pivot irrigation were sampled in southern Georgia to assess chemical differences between non-irrigated and irrigated areas. Proton (^1^H) nuclear magnetic resonance (NMR) data were obtained from cotton leaves, and Principal Component Analysis (PCA) was performed to assess differences in chemical composition. Across all samples, farm site accounted for most of the variability, but within each farm site, the PCA scores plots showed clear separation between non-irrigated and irrigated conditions in 10 sites. Inspecting the PCA loadings revealed significant resonances resembling a lipid-like signal. After reverse-phase fractionation, we observed that many of these resonances appeared together in later fractions, suggesting a lipid, specifically a fatty acid such as linoleic acid. We hypothesized that differences in net lipid saturation level may drive separation between non-irrigated and irrigated samples. Six farm sites had a significantly or marginally significantly higher degree of unsaturation in irrigated samples, while one farm site had significantly higher unsaturation in non-irrigated samples. Our results indicate that drought stress likely affects lipid profile composition, which could be driving higher herbivorous pest densities in drought-stressed crops.

## Introduction

Cotton (*Gossypium hirsutum*) is a vital textile fiber and oilseed crop grown worldwide. It is more tolerant of abiotic stressors than most plants. Still, drought stress can affect plant height, leaf area, dry weight, fiber quality, and root development, and even reduce cotton leaves’ photosynthetic rate, transpiration rate, and stomatal conductance (Zhang et al., 2021). In southern Georgia, approximately 1.1 million acres are used to grow cotton annually, with at least 45% of the crop irrigated using center-pivot systems. Despite extensive irrigation infrastructure, plants farther from the centers of these systems may be more exposed to heat and water stress. Furthermore, inadequate irrigation can reduce cotton’s ability to defend against herbivores, such as insect pests. This results in drought-stressed plants that are chemically more nutritious, less toxic, or both to insect herbivores.

Increases in global temperatures paired with changing precipitation patterns have exacerbated outbreaks of agricultural insect pests by advancing insect phenology, allowing distribution/dispersal to new areas, and speeding physiology and development (Bale et al., 2002; Bannerman, Gillespie, & Roitberg, 2011; Chen, 2016; Deutsch et al., 2018; Gutierrez, Ponti, d’Oultremont, & Ellis, 2008). The silverleaf whitefly (*Bemisia tabaci*) has recently become a dominant pest of field crops in the southeastern US, and the effects of climate change-associated drought may have led to the sudden transition to major pest status. For example, climate change’s high temperatures and water stress can likely shorten the development time of multivoltine species, leading to more pest outbreaks in crops such as cotton (Aslam, Johnson, & Karley, 2013; Bale et al., 2002). These outbreaks often lead to lower crop yield (Khizar et al., 2020).

Most plants, including cotton, have chemical defenses to protect themselves against pests such as the silverleaf whitefly (A. Mithöfer & Boland, 2012). Some general chemical defensive compound classes in plants include alkaloids, terpenoids, phenolics, glucosinolates, flavonoids, anthocyanins, and polypeptides (Axel Mithöfer & Maffei, 2016; Satish et al., 2020). Beyond defense, many of these compounds also accumulate in response to abiotic stresses. Drought can also significantly alter metabolic processes, thereby altering the chemical composition of defense-related metabolites (Khan et al., 2025). Profiling changes in the cotton metabolome under stress and non-stress conditions can reveal important metabolite pathways, and this may help us better understand how to protect against pests.

To investigate changes in the metabolome of cotton as a response to drought and poor irrigation, we sampled cotton leaves in irrigated and non-irrigated conditions from 20 different farms in southern Georgia. In this study, we used Nuclear Magnetic Resonance (NMR) spectroscopy because it can detect and quantify metabolites in complex matrices, including raw cotton leaf extracts, without the need for selective targeting (Dillon et al., 2024). NMR is highly reproducible, enabling robust interpretation of high-dimensional data through methods like principal component analysis in untargeted metabolic profiling (Edison et al., 2021). We hypothesized that drought changes the chemical profile of the cotton endometabolome compared to irrigated conditions.

## Materials and Methods

### Sample collection and preparation

Cotton (*Gossypium hirsutum*) leaves were collected from 20 variously managed cotton farms in southern Georgia, all of which used a center-pivot irrigation system (Figure 1). The non-irrigated corners resulting from center-pivot irrigation provided us with a unique opportunity to leverage irrigated and non-irrigated treatments within each farm site (Figure S1). Five leaves from individual plants were combined into 50 mL Falcon tubes, flash-frozen with liquid nitrogen, and kept on dry ice before storage at –80 °C. In total, 120 cotton leaf samples (20 farms x 2 treatments x 3 replicates) were collected. All samples were randomized, and quality control samples were added to compose the analytical sequence.

**Figure 1:**
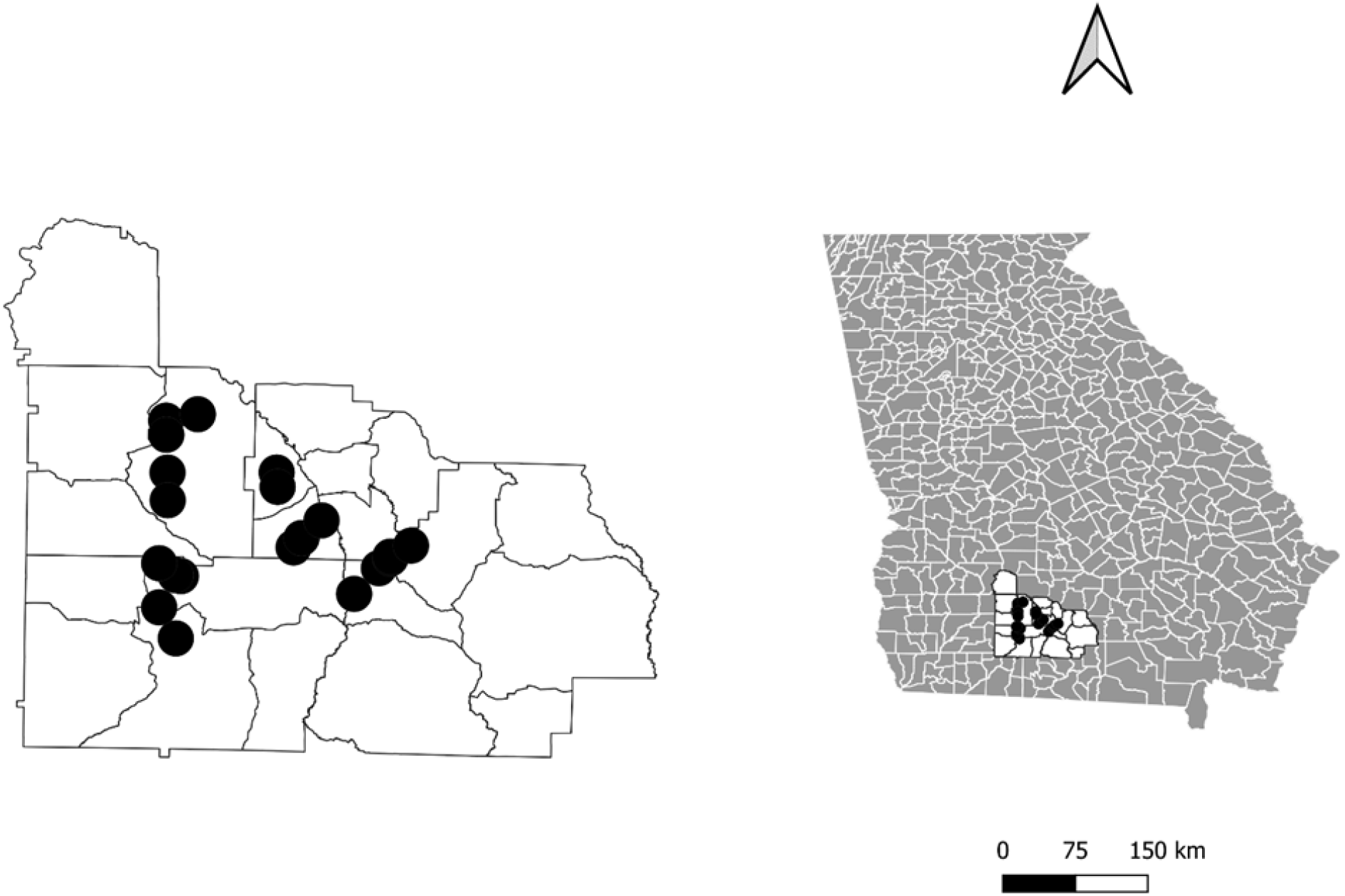
Sampled Farm Sites in South Georgia

### Sample extraction

The samples were extracted according to this protocol: (dx.doi.org/10.17504/protocols.io.dm6gpjwkdgzp/v2). In short, the samples were taken from the freezer, lyophilized for 24 hours, and then macerated into a powder. For each sample, 50 mg of dry plant material was weighed and transferred to 2 mL screw cap microtubes. An equivalent of 200 µL of 1 mm Zirconia beads and 1.5 mL of methanol were added to each tube. The samples were homogenized using a bead beater and then centrifuged at 14,000 rpm for 15 min. After centrifugation, 750 µL of the supernatant was transferred to separate tubes. An additional 750 µL of methanol was added to the original tubes, and the pellets were reconstituted via vortexing and sonication. The samples were again centrifuged at 14,000 rpm for 15 min, and 750 µL of the supernatant was transferred to the same tubes as the previous transfer. Pooled samples were then created, and all the samples were dried in a Centrivap and stored at −20 °C.

### One-dimensional (1D) NMR

Samples were reconstituted in 60 μL of Deuterated Dimethylsulfoxide (DMSO-D6) with 0.03% TMS (internal standard) and transferred to 1.7 mm Bruker SampleJet NMR tubes. All NMR data were collected using a Bruker Avance Neo console on an Oxford 800 MHz magnet equipped with a 1.7 mm TCI cryoprobe and SampleJet sample changer. One-dimensional NMR data were acquired at 310K using a “noesypr1d” pulse sequence, and 32,768 points were collected with 4 dummy scans and 32 scans for each sample. An exponential window function of 0.3, Fourier Transform, automatic phasing, and referencing were done using the NMRPipe batch processing scheme (dx.doi.org/10.17504/protocols.io.kqdg3xen1g25/v1). Data were imported into MATLAB, and alignment using Constrained Correlation Optimized Warping (CCOW) was done using the Edison Lab Metabolomics Toolbox (https://github.com/edisonomics/metabolomics_toolbox). The aligned data were then uploaded to the MATLAB software PLS_Toolbox and preprocessed by removing end and solvent regions, PQN normalization, mean centering, and Pareto scaling before PCA analysis (PLS_Toolbox 9.3 (2023) Eigenvector Research, Inc., Manson, WA USA 98831; software available at http://www.eigenvector.com).

### Two-dimensional (2D) NMR

Two-dimensional (2D) NMR data were collected on fraction 140, which contained high-loading peaks from PCA. HSQC and TOCSY spectra were acquired at 298K. The HSQC spectrum was acquired with 8 scans, t1 and t2 acquisition times of 7.7 and 188.4 ms, and data points in f1 and f2 of 512 and 4096, respectively.

The HSQC was also collected with a sweep width of 13.59 ppm in the direct dimension and 165 ppm in the indirect dimension. The TOCSY spectrum was acquired with 8 scans, t1 and t2 acquisition times of 23.6 and 188.4 ms, and data points in f1 and f2 of 512 and 4096, respectively. The TOCSY was collected with a sweep width of 13.59 in both direct and indirect dimensions and a mixing time of 80 ms.

Both 1D and 2D raw data are available on the NAN resource connector (Hoch et al. 2025).

### Reverse Phase Fractionation

A pooled QC sample was reconstituted in 100% methanol. Seven injections of 100 µL were fractionated using an Agilent 1260 Infinity HPLC with Avantor ACE C18-AR, 100 Å, 5 µm, 10 × 150 mm HPLC column at 40 °C. A 35 min gradient of 0.05 % formic acid in H_2_O, 0.05% formic acid in ACN, and Isopropanol was used for the fractionation (Table S1). From 3 to 14 min, 120 equally spaced fractions were collected, and from 19.45 to 28 min, 20 equally spaced fractions were collected. The fractions were dried in a Centrivap at room temperature, then stored at −80 °C until NMR analysis.

### Calculation of Lipid Degree of Unsaturation

The degree of unsaturation (*fu* Index) was calculated by taking the ratio of the integrated signal of allylic protons to that of α-methylene protons, as described in Mosconi et al. (2011) and Sadeghi-Nik (2026). Because the samples were collected in DMSO, the chemical shifts of the signals are shifted compared to those collected in chloroform. The allylic signals were integrated from 1.921 – 1.991 ppm, and the α-methylene signals were integrated from 2.176 – 2.208 ppm. After integration, the degree of unsaturation was calculated (Eq. 1), and a two-sample t-test was done between the *fu* Index of irrigated and non-irrigated samples for each farm site and between all farm sites.

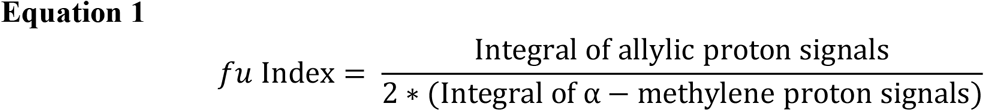

## Results

### Multivariate Analysis Reveals Variation between Farm Sites

We first used Principal Component Analysis (PCA) to investigate differences between irrigated and non-irrigated conditions across all 20 farm sites (Figure S2). However, the analysis revealed minimal separation between the two conditions. We speculated that site-specific variability outweighed the effects of irrigation differences; for example, confounding factors, such as elevation shifts, may have allowed samples in the non-irrigated condition to receive unintended moisture. Consequently, we excluded 10 farms whose PCA score plots failed to demonstrate a clear distinction between irrigated and non-irrigated conditions within the farm site (Figure S3).

Using the 10 included farm sites, we again used PCA to explore variation between irrigated and non-irrigated conditions (Figure 2). The scores plot revealed a slight separation between irrigated and non-irrigated conditions, but it was not strongly defined. With further inspection, the PCA scores plots colored by farm site indicated that the primary source of variation remained between farm sites rather than between the irrigated and non-irrigated treatment conditions. This suggests that site-specific factors may contribute more significantly to the observed metabolic differences than irrigation conditions.

**Figure 2:**
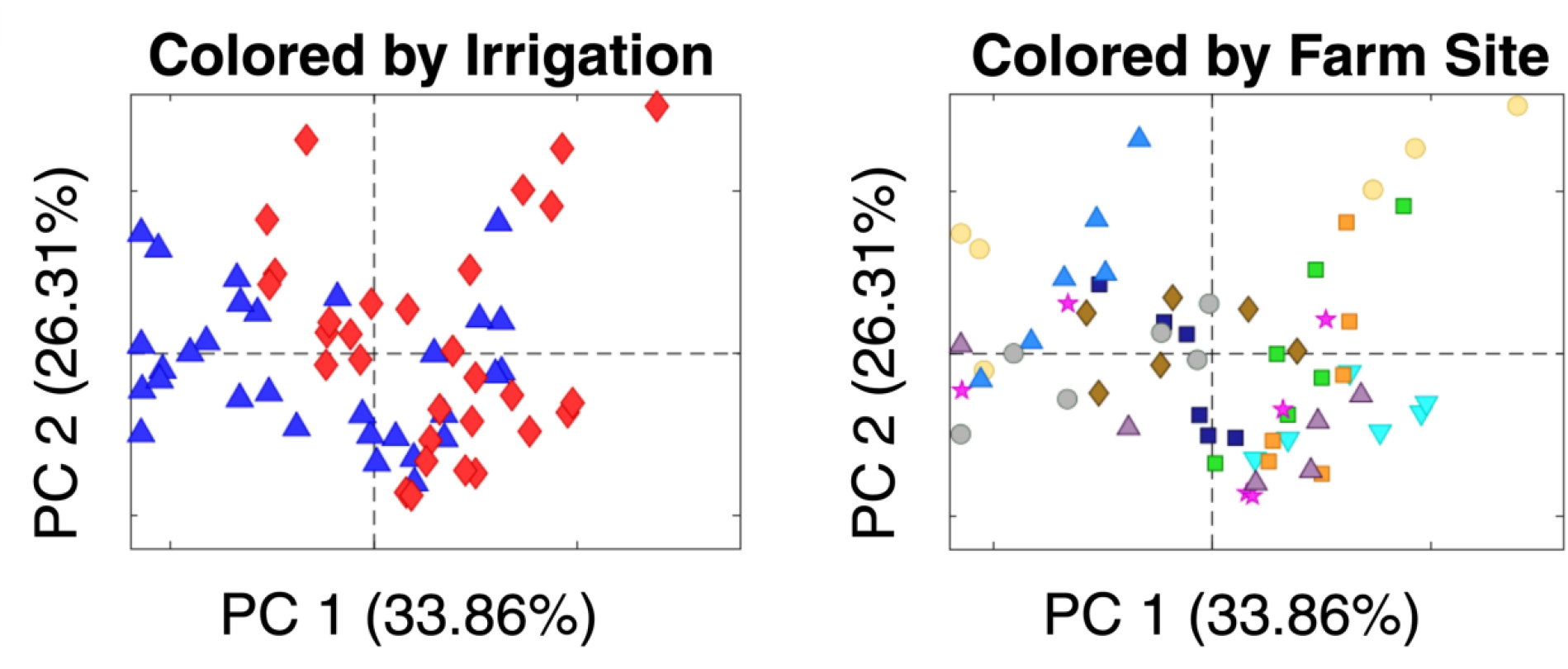
**a**, Principal Components Analysis scores plot colored by irrigated (blue) vs non-irrigated (red) condition. **b**, The same scores plot colored by farm site.

### Differences in Irrigated and Non-irrigated Cotton Metabolome Within Farm-Sites

To further investigate the cotton metabolome under irrigated versus non-irrigated conditions, we analyzed PCA scores plots site-by-site (Figure 3). Clear separation between irrigated and non-irrigated conditions was observed at each farm site, indicating that irrigation status impacts the metabolic profiles of cotton plants at a local level. This separation suggests that, while overarching farm-site differences play a larger role in overall metabolic variability, irrigation status still contributes to discernible changes in chemical composition at the within-site level.

**Figure 3:**
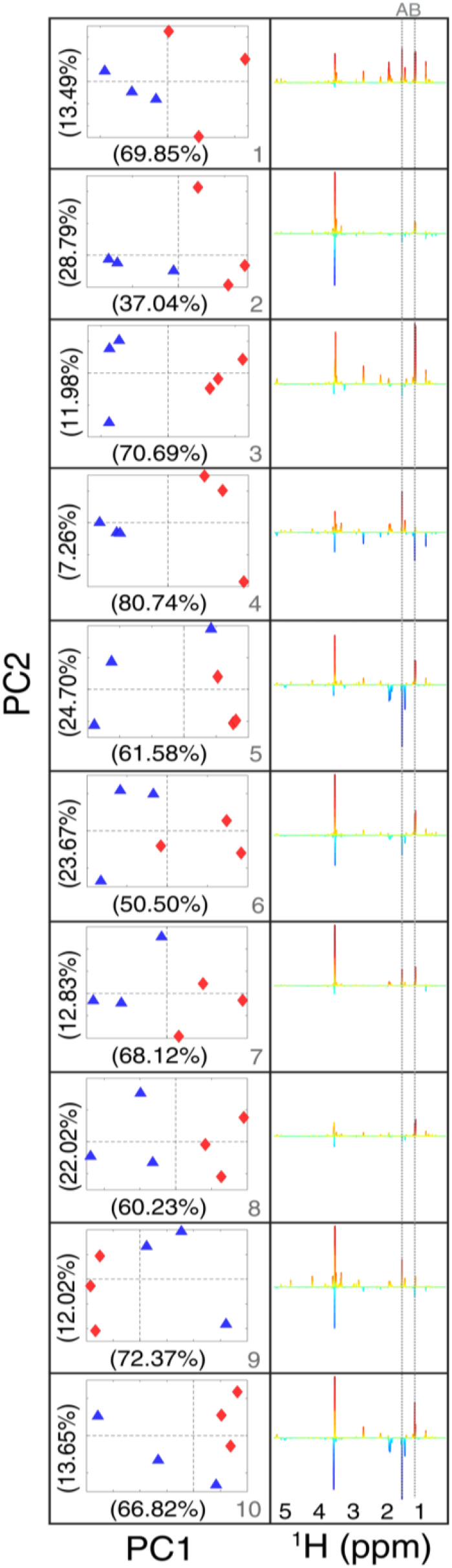
Principal Components Analysis scores and loadings plots of 10 individual farm sites. The scores plots are colored by irrigated (blue) vs non-irrigated (red). The loadings are along the first principal component. The gray dotted lines labeled A & B indicate resonances shared between many of the farm sites.

### Similar Chemical Species in Loadings

To identify the key signals driving separation in the PCA scores plots, we analyzed the loadings along the first principal component from the within-farm PCA models (Figure 3). Visual inspection revealed several patterns; for instance, a peak appearing at 1.16 ppm (peak B, labeled with the gray dotted line across each plot) consistently associated with the non-irrigated condition (red positive peaks). Interestingly, in Farm 4, this same peak appeared to be associated with the irrigated condition (blue negative peaks). We also observed several peaks at 1.56 ppm with large loadings in the irrigated condition, shared between Farms 5 and 10 (peak A). We then quantitatively analyzed the loadings to determine which chemical shifts ranked most consistently among the top 10 absolute loading scores for each site (Figure S4). Both visually identified peaks A and B were indeed the most common drivers of separation.

### High Loadings Peaks Elute in the Same Fraction

Because of the spectral complexity of our sample, we performed reverse-phase chromatography on a pooled mixture and collected one-dimensional (1D) NMR data on each of the 140 fractions. By looking at our most frequent loadings, we noticed that many of the top peaks appear in the last fractions of our separation (Figure 4). This led us to believe these resonances may be part of the same molecule, and it is most likely non-polar as it eluted at the end of the reverse-phase chromatography. We collected HSQC and TOCSY spectra on fraction 140 to help us identify the molecule. We uploaded the 2D data to the web server COLMAR for database matching. We matched against the COLMAR non-polar database and found the closest match to the lipid linoleic acid. To confirm whether linoleic acid was the molecule, we spiked a linoleic acid standard into fraction 140 and recollected 1D NMR data (Figure S5). We observed new peaks appear in the fraction, indicating that linoleic acid was not the correct compound.

**Figure 4:**
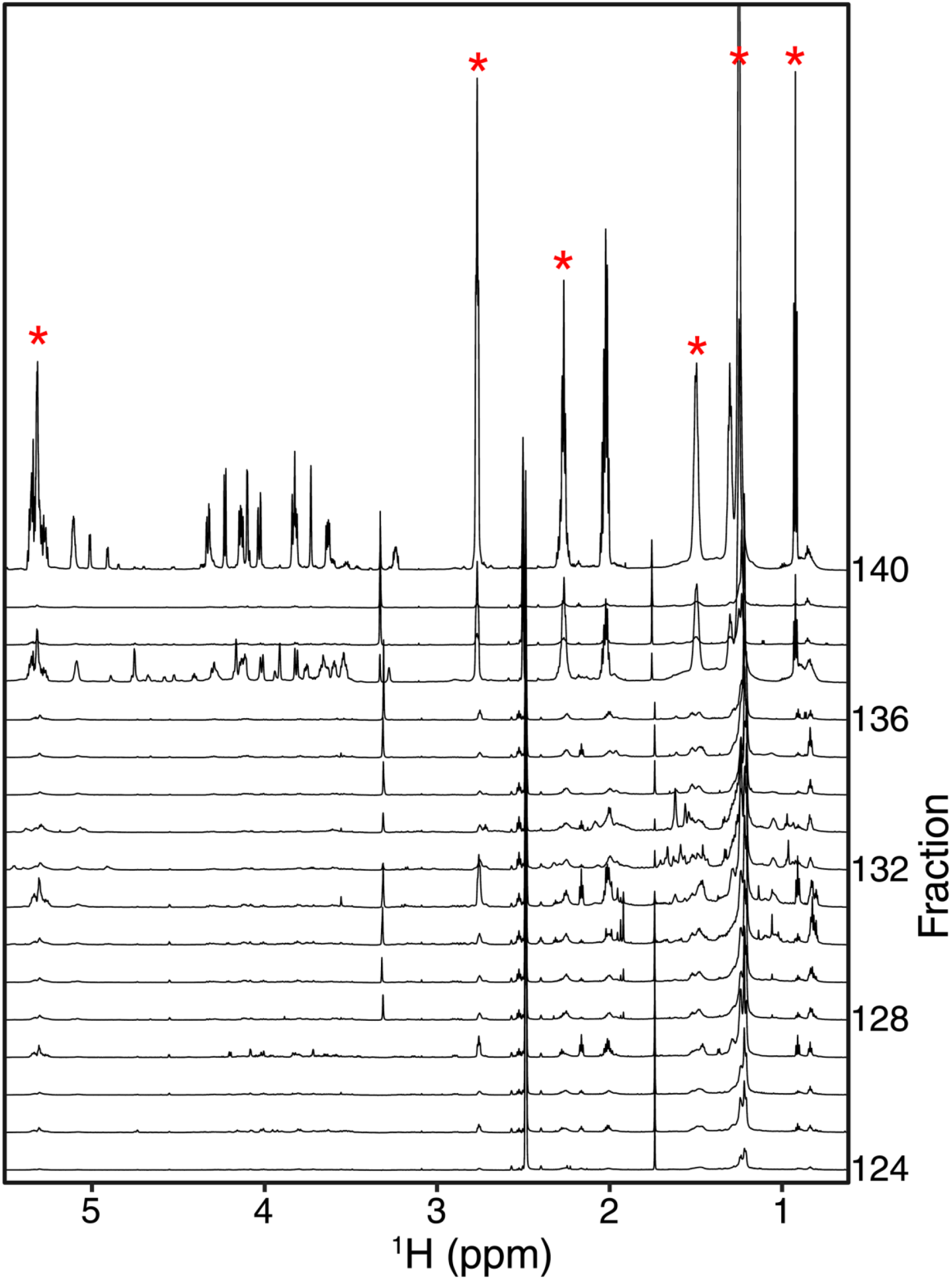
Last 16 fractions from a reverse phase separation of cotton leaf extracts. Peaks found to be important in the loadings are highlighted with a red asterisk.

### Differences in Lipid Saturation between farm sites

Lipids are dominated by characteristic methylene and other aliphatic NMR signals, making annotation difficult. We next examined whether the overall level of unsaturation differed between the irrigated and non-irrigated groups. We calculated the degree of unsaturation within each farm site and then performed a two-sample t-test for each group. To calculate the degree of unsaturation (*fu* Index), we calculated the ratio of allylic and α-methylene proton signals in the samples as described in Mosconi et al. (2011) and Sadeghi-Nik (2026) (Eq. 1). Based on our calculation of unsaturation levels, we observed a trend: the irrigated group tended to have higher unsaturation levels (Figure 5). At farm site 4, however, the degree of unsaturation was significantly higher in the non-irrigated group than in the irrigated group. We also calculated the *fu* Index across all sites and found that the irrigated samples had a significantly higher degree of unsaturation than the non-irrigated samples (Figure S6a). Interestingly, the degree of unsaturation between the ten farm sites that showed the least separation in PCA scores between irrigated and non-irrigated conditions was not significantly different, indicating that the degree of unsaturation was important for separation between irrigated and non-irrigated plants (Figure S6b).

**Figure 5:**
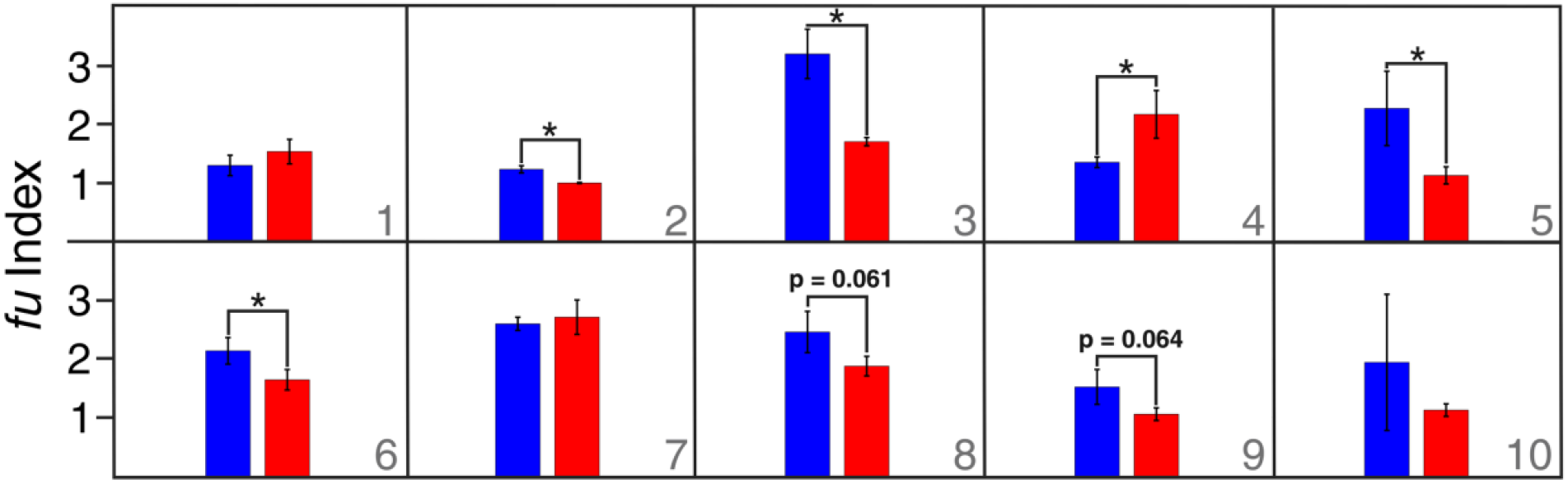
Unsaturation levels of lipids of irrigated vs non-irrigated cotton leaves within individual farm sites.

## Discussion

Our analysis demonstrated metabolic differences between irrigated and non-irrigated cotton across several farms in southern Georgia. These metabolic differences were primarily driven by the degree of leaf lipid saturation. Due to variability across farm sites, these differences were observed only within sites and only for a subset of the sites. It remains unclear what is causing the difference in lipid profiles, but it has previously been reported that abiotic stresses induce specific effects on plant membranes. Similarly, it is unclear whether these changes affect the nutritional quality of the foliage or overall plant chemical defense from insect herbivores.

Membrane lipid remodeling is defined as the alteration of fatty acid composition and the change in degree of saturation in fatty acids (Liu et al. 2019). Plants are known to alter the fatty acid composition of membrane lipids to improve stress tolerance under drought, heat, and cold. (Ou et al. 2025; Liu et al. 2019). For example, a recent study examined heat-tolerant and heat-sensitive wheat genotypes and found that they exhibited varying degrees of changes in leaf lipid composition and fatty acid unsaturation (Narayanan et al. 2016). Similar to our findings, other studies indicate that the degree of unsaturation decreased under drought stress (Dakhma et al. 1995; Zhang et al. 2005; Xu et al. 2011). For example, in safflower, C18:2 and C18:3 contents were reduced (Hamrouni et al. 2001), and in sage, there was a decrease in 16:1 and 18:3 fatty acids (Bettaieb et al. 2009). Meta-analysis revealed reductions in levels of 16:1, 16:3, and 18:3 fatty acids, suggesting that membrane fatty acid unsaturation was reduced upon experiencing stress such as drought (Liu et al. 2019). When examining the drought tolerance of Kentucky bluegrass cultivars, the alteration in fatty acid unsaturation level was found to be induced by drought via changes in the composition of linolenic (18:3), linoleic (18:2), palmitic (16:0), and stearic acids (18:0) (Xu et al. 2011). It appeared that fatty acids became more saturated as relative water content (cellular hydration) declined, but this depended on the cultivar, with better membrane stability and maintenance of higher fatty acid unsaturation levels. In tobacco plants, drought was found to reduce linolenic acid levels, indicating damage (Zhang et al. 2005). On the other hand, when investigating the drought tolerance of the coconut palm, researchers found that linolenic acid was enriched in response to water deficit despite the overall reduction in lipid contents (Repellin et al. 1997). Linoleic acid is increasingly being reported to be modulated in response to stressors such as drought, low temperature, and high salinity via alterations in the membrane fluidity and stability (Liu et al. 2019; Ou et al. 2025). Studies investigating lipid remodeling in response to drought have been limited to crops such as rice, wheat, bean, cowpea, and maize (Kaoua et al. 2006; Torres-Franklin et al. 2007; Martins Júnior et al. 2008; Liu et al. 2011; Zi et al. 2022; Yin et al. 2024).

The plant membrane is significant in photosynthesis and can be regulated to improve cellular metabolism and reduce physiochemical damage caused by drought (Yin et al. 2024). Water stress or drought stress can dismantle essential phospholipids in the membrane and increase the production of signaling lipids while decreasing essential oil production (Sharma et al. 2023; Henschel et al. 2024). Other ways plants can regulate the synthesis and composition of several lipid classes include increased cuticle deposition of cutin and waxes, which may help reduce transpiration by sealing the leaf surface (Sharma et al. 2023; Henschel et al. 2024). Plant surface waxy compounds include long-chain alkanes, alcohols, carboxylic acids, and secondary metabolites such as quinones and flavonoids (Ali et al. 2021). Drought stress may exacerbate silverleaf whitefly (SLWF) outbreaks (*Bemisia tabaci*) by increasing nutritional quality or decreasing defenses. The nutritional quality of the cotton leaves may be greatly affected by water and heat stress, thus attracting greater numbers of whiteflies. If that were the case, the heat and water stress may have caused the plants to allocate resources to nutrition and growth rather than defenses against herbivore attacks. When insects such as whiteflies and aphids approach a plant to attack, they are first faced with cuticular compounds, such as 1-hexacosanol, short-chain fatty acids, carboxylic acids, and wax esters with antixenotic properties to discourage insects from attacking and feeding (Ali et al. 2021). Since no variety of *Gossypium hirsutum* has been reported as resistant against the cotton leaf curl disease, a study examined the epicuticular waxes of different cotton genotypes since the increased amounts of epicuticular waxes have been deemed as the leading barriers for whiteflies in *Gossypium arboreum* (Ali et al. 2021). The researchers found that *G. hirsutum* was more susceptible to whitefly visits and showed the presence of five compounds: trichloroacetic acid, hexadecylester, P-xylenolphtalein, 2-cyclopentene-1-ol, 1-phenyl-, and Phenol, 2.5-bis [1,1-dimethyl], which showed a bonding within the chitin of the whitefly. Though the exact mechanisms of insect interaction and repulsion are not clear, the amount and composition of wax play clear roles in attracting and repelling whiteflies.

Cotton has also been found to produce foliar terpenoid aldehydes, including gossypol, hemigossypolone, and heliocides, which, in response to physical wounding, increased in younger leaves, while concentrated carbohydrates, amino acids, and fatty acids accumulated in the nectar (Park et al. 2019), which attracts both damaging and beneficial pests. Interestingly, little is known about how plant membrane fluidity and stability can be physiologically overcome by piercing-sucking herbivores such as aphids and whiteflies, and whether this is made easier or harder by abiotic stresses such as drought.

Metabolomic analysis of plants by NMR spectroscopy allowed us to robustly profile many plants from across southern Georgia farms. We used this profiling data to identify important peaks causing separation between our samples; however, due to the complexity of the plant metabolome and the lack of database spectra for compounds collected in DMSO, we were unable to identify the specific molecules these peaks corresponded to. We selected DMSO to maximize the solubility of secondary and hydrophobic metabolites in our extractions. No single analytical technology can cover the entire plant metabolome; thus, different extraction techniques and complementary analytical methods, such as gas chromatography-mass spectrometry (GC-MS) and NMR, are often used. Because NMR is inherently quantitative, we were able to assess lipid unsaturation levels from the spectra.

Drought stress may exacerbate silverleaf whitefly (SLWF) outbreaks, *B. tabaci*, by increasing nutritional quality or decreasing defenses. The nutritional quality of the cotton leaves may be greatly affected by water and heat stress, thus attracting greater numbers of whiteflies. If that were the case, the heat and water stress may have caused the plants to allocate resources to nutrition and growth rather than defenses against herbivore attacks. The regional spraying of pesticides to control whiteflies, in conjunction with warmer temperatures (degree-days), resulted in greater outbreaks of SLWF (Crossley et al. 2023). Whiteflies are reported to exhibit resistance to several major classes of insecticides, including over 60 active ingredients (Denholm et al.1998; Horowitz et al. 2020; Perier et al. 2022). The ongoing evolution of resistance in the SLWF enables it to evade host plant defenses and artificial chemical control, leading to greater plant and crop damage. It is worth investigating further how defensive chemicals, membrane fluidity, and piercing-sucking herbivory relate to bottom-up mechanisms that may be exacerbating pest outbreaks in crops. These ecological mechanisms can now be investigated at the molecular level to generate “ecometabolomic” studies (Sardans et al. 2020) that better enable us to understand organisms’ responses in a drastically changing environment.

## Acknowledgements

ASE and CE were supported by NIH R35GM148250.

This work was funded by the USDA-Agricultural Research Service Non-Assistance Cooperative Agreement #58-6080-9-006 “Managing whiteflies and whitefly-transmitted viruses in vegetable crops in the southeastern U.S.”

## Data Availability

Both 1D and 2D raw data and metadata are available on the NAN resource connector (Hoch et al. 2025).

## Supplement

**Figure S1.**
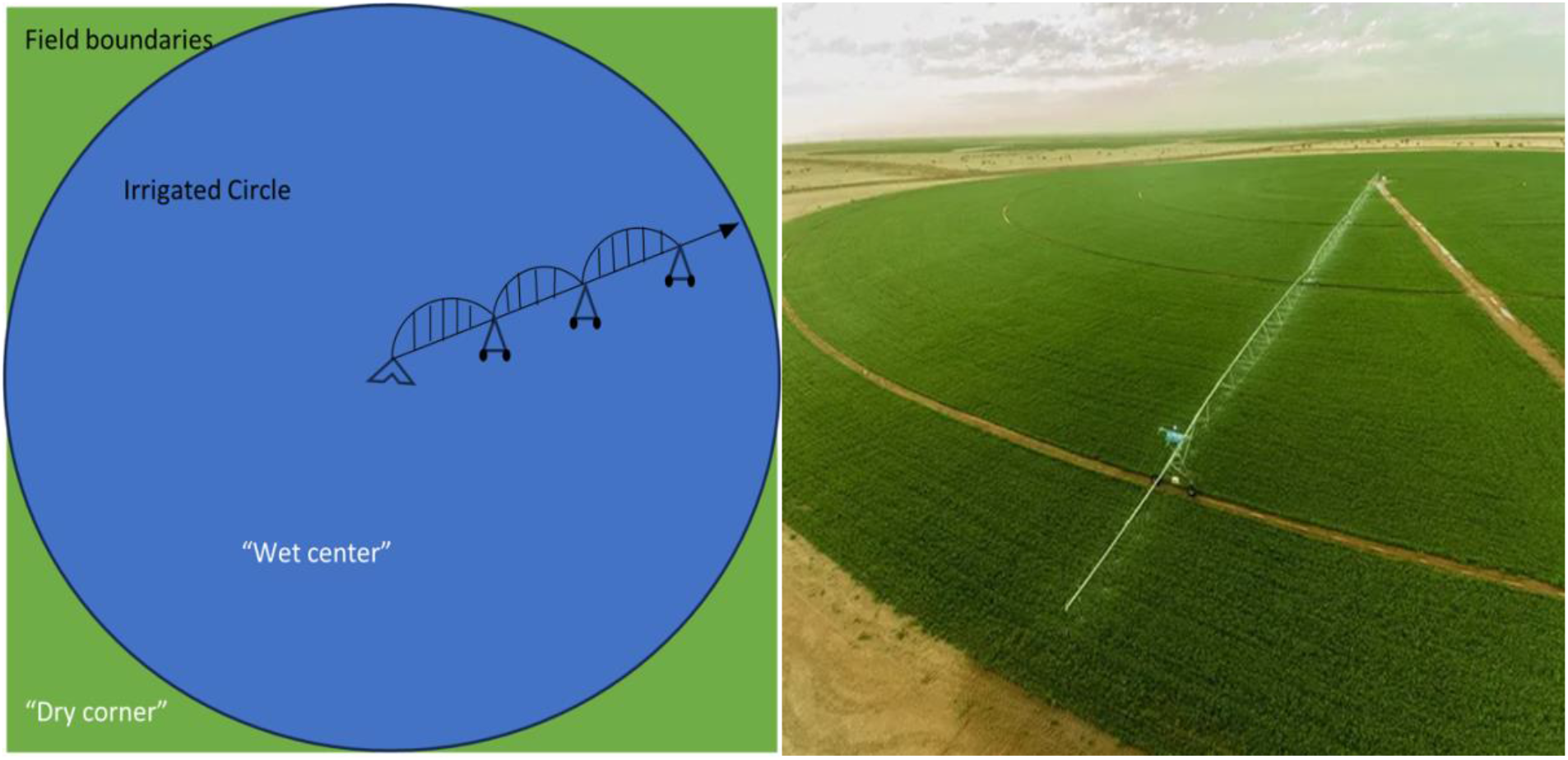
Center-pivot irrigation system in a square field resulting in a circle of irrigated (wet) crops and corners of drought-stressed (dry) crops; image (right) by AFKO - Small Center Pivot Irrigation System by AFKO Ltd. Şti.

**Table S1.** Gradient for Reverse Phase Fractionation. A 35-minute gradient of 0.05 % formic acid in H_2_O (A), 0.05% formic acid in ACN (B), and Isopropanol (C) was used for the fractionation

| Time (Min) | A (%) | B (%) | C (%) | Flow (mL/min) |
| --- | --- | --- | --- | --- |
| 0.0 | 95.0 | 5.0 | 0.0 | 2.0 |
| 2.0 | 90.0 | 10.0 | 0.0 | 2.0 |
| 4.0 | 70.0 | 30.0 | 0.0 | 2.0 |
| 5.0 | 67.0 | 33.0 | 0.0 | 2.0 |
| 15.0 | 57.0 | 43.0 | 0.0 | 2.0 |
| 18.0 | 20.0 | 80.0 | 0.0 | 2.0 |
| 21.0 | 2.0 | 98.0 | 0.0 | 2.5 |
| 26.0 | 2.0 | 98.0 | 0.0 | 2.5 |
| 27.0 | 0.0 | 80.0 | 20.0 | 2.0 |
| 30.0 | 0.0 | 80.0 | 20.0 | 2.0 |
| 30.5 | 95.0 | 5.0 | 0.0 | 2.0 |
| 35.0 | 95.0 | 5.0 | 0.0 | 2.0 |

**Figure S2.**
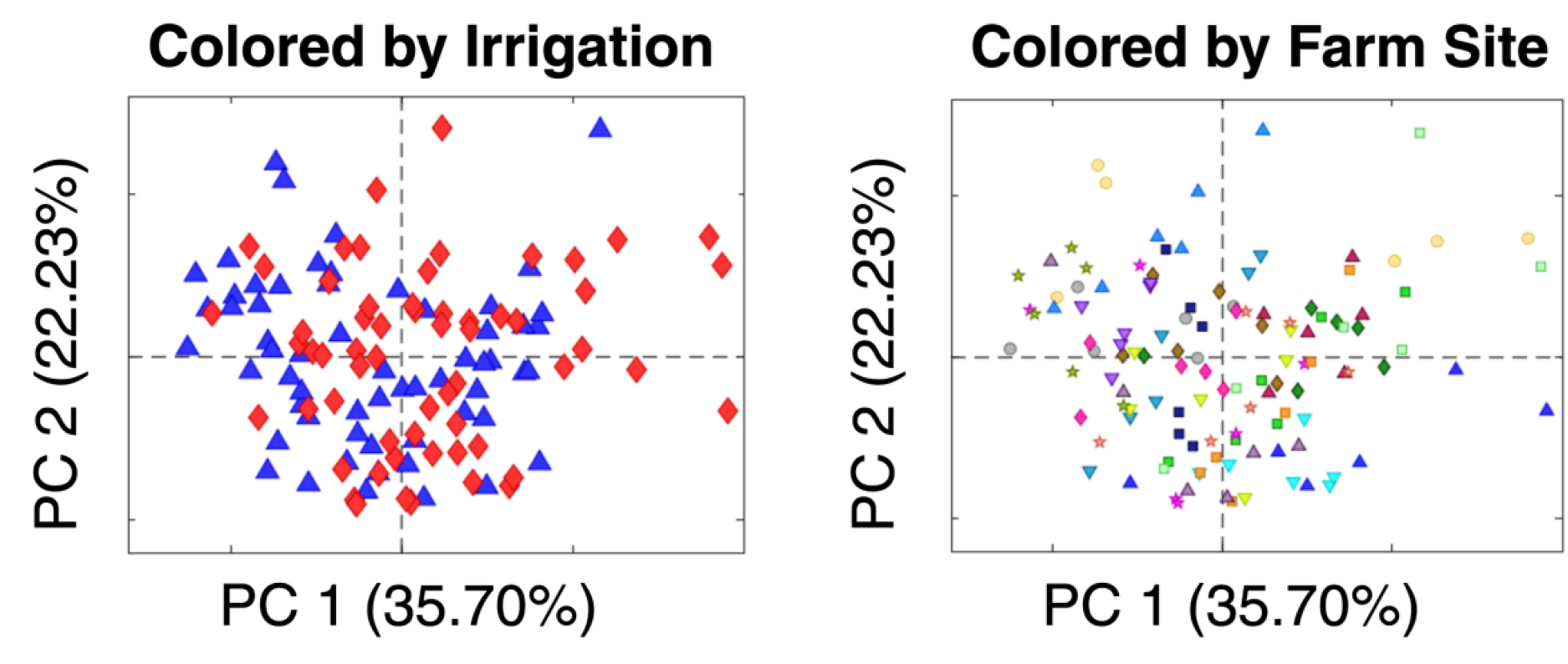
PCA scores plots of all 20 farm sites. The first plot is colored by irrigated (blue) and non-irrigated (red) condition, and the second plot is colored by farm site.

**Figure S3.**
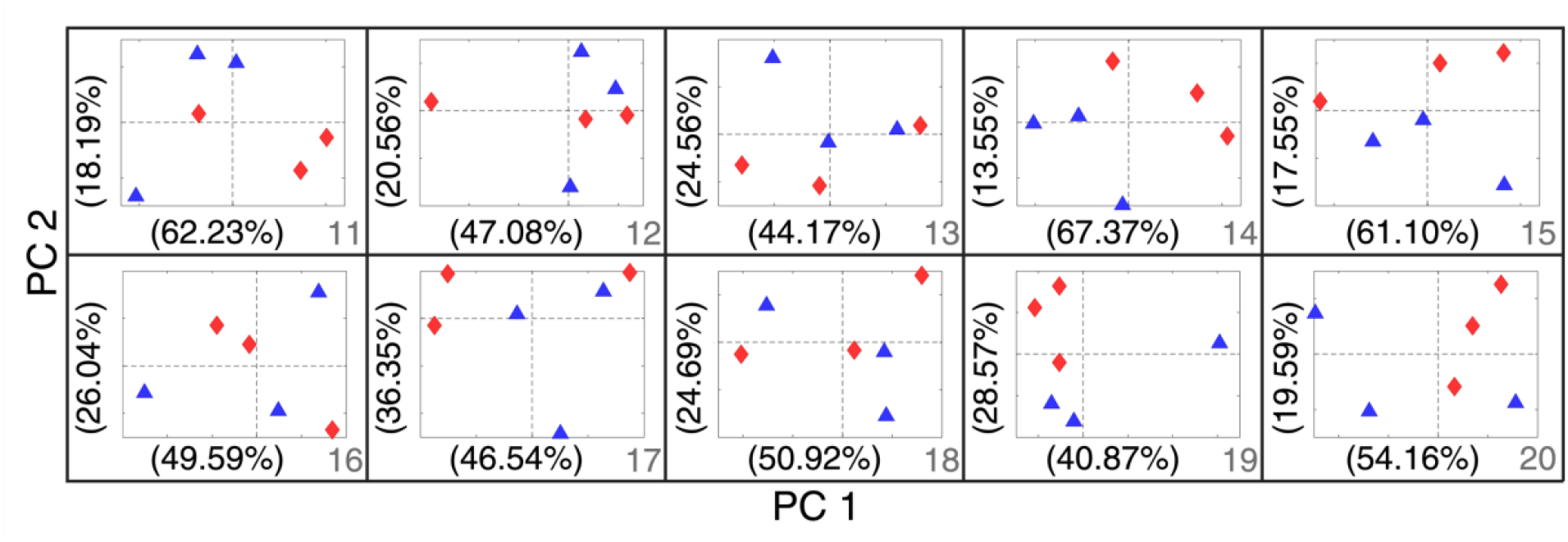
PCA scores plots of 10 farms showing less separation between irrigated (blue) versus non-irrigated (red) conditions.

**Figure S4.**
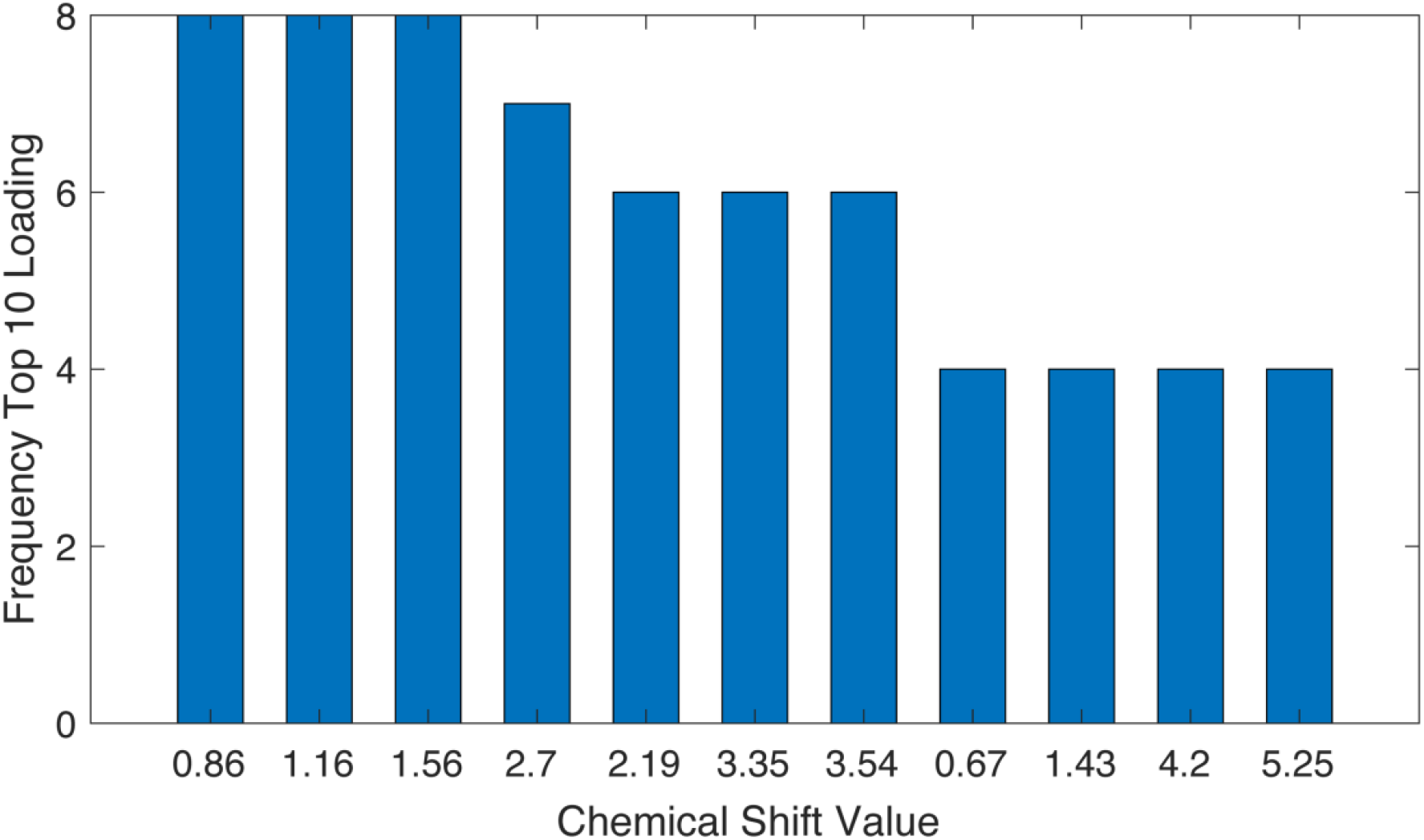
Frequency of peaks that appear in the top 10 absolute loadings.

**Figure S5.**
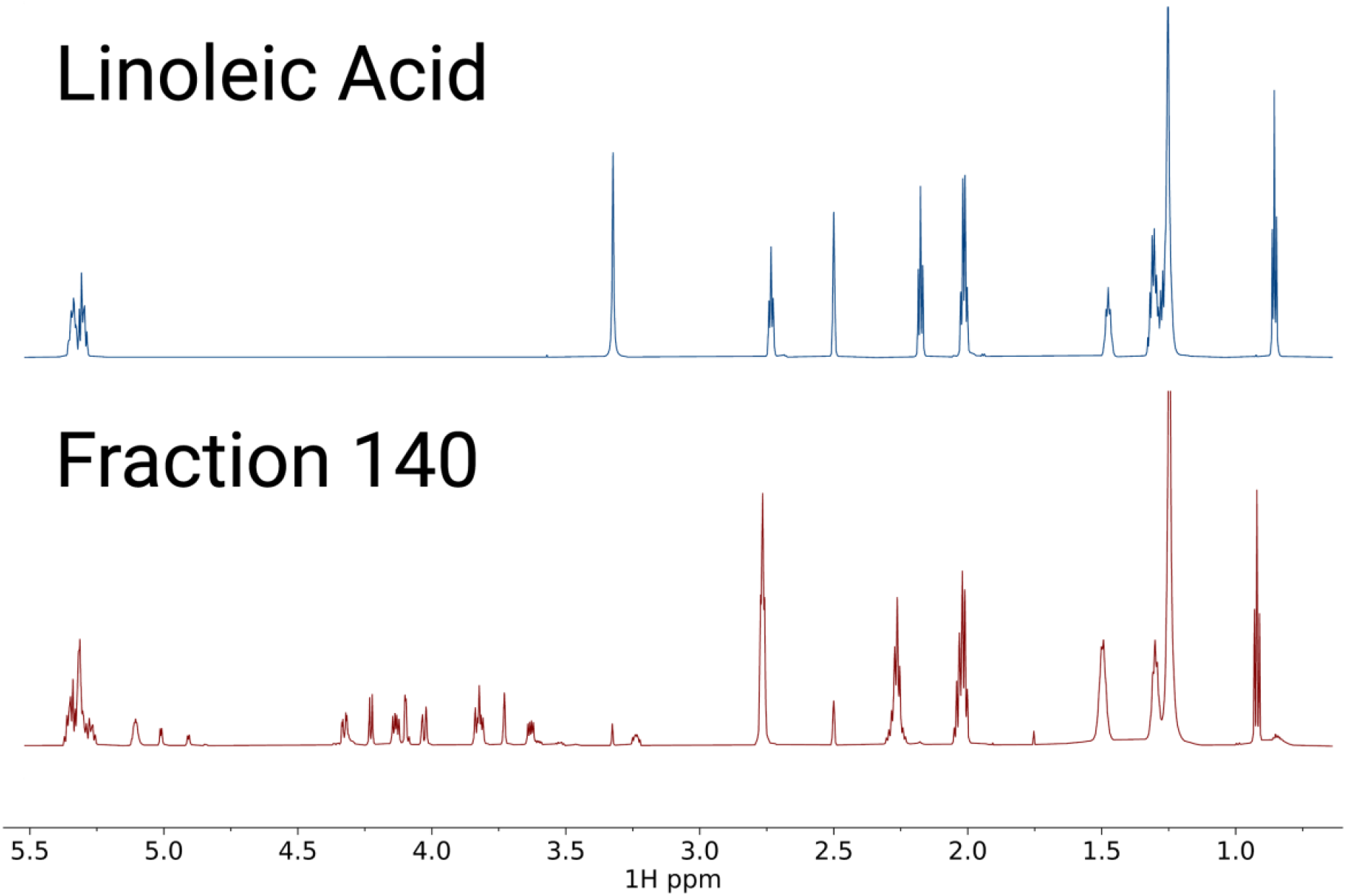
Linoleic Acid standard spectrum versus fraction 140.

**Figure S6.**
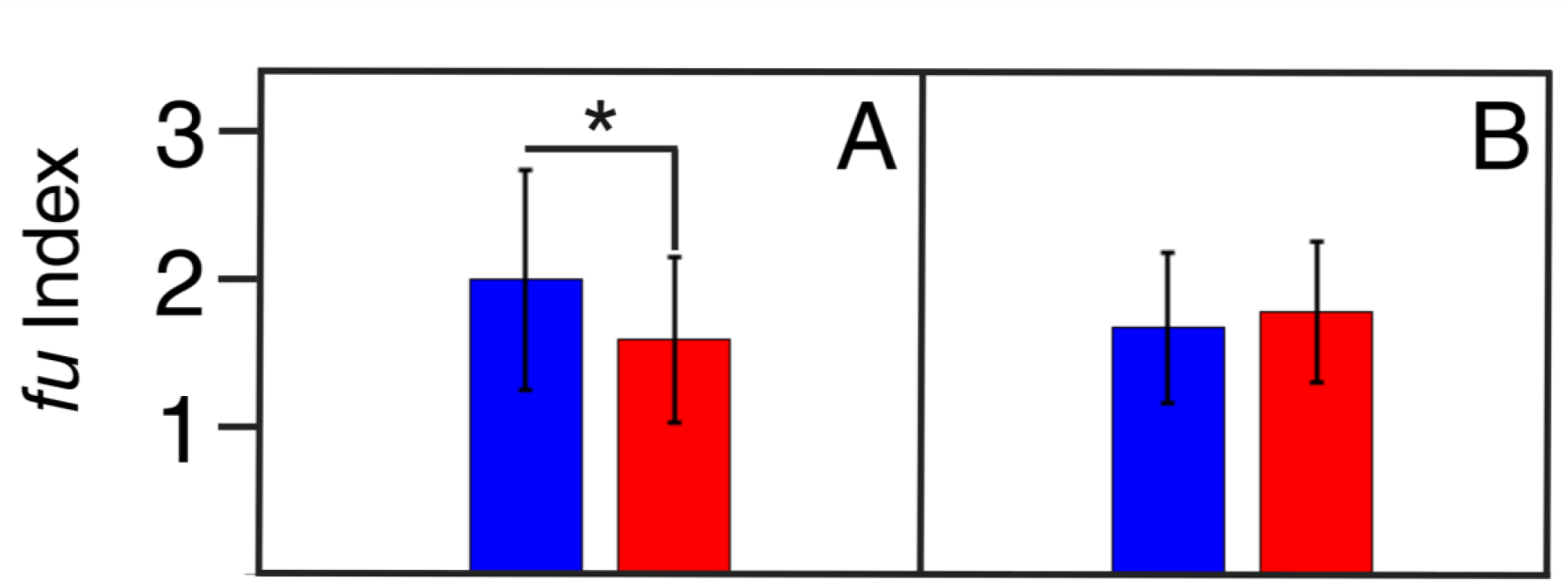
T-test of the degree of unsaturation between all farm sites. (A) The ten farms that showed the greatest separation in PCA scores between irrigated and non-irrigated samples have a significant difference in the degree of unsaturation between the two conditions. (B) The ten farms with little difference between wet and dry in the PCA scores do not show a significant difference in the degree of unsaturation between the two conditions.

